# Kinetochore protein Spindly controls microtubule polarity in *Drosophila* axons

**DOI:** 10.1101/2020.03.20.000364

**Authors:** Urko del Castillo, Hans-Arno J. Müller, Vladimir I. Gelfand

## Abstract

Microtubule polarity in axons and dendrites defines the direction of intracellular transport in neurons. Axons contain arrays of uniformly polarized microtubules with plus-ends facing the tips of the processes (plus-end-out), while dendrites contain microtubules with minus-end-out orientation. It has been shown that cytoplasmic dynein, targeted to cortical actin, removes minus-end-out microtubules from axons. Here we have identified Spindly, a protein known for recruitment of dynein to kinetochores in mitosis, as a key factor required for dynein-dependent microtubule sorting in axons of *Drosophila* neurons. Depletion of Spindly affects polarity of axonal microtubules *in vivo* and in primary neuronal cultures. In addition to these defects, depletion of Spindly in neurons causes major collapse of axonal patterning in the third-instar larval brain as well as severe coordination impairment in adult flies. These defects can be fully rescued by full-length Spindly, but not by variants with mutations in its dynein-binding site. Biochemical analysis demonstrated that Spindly binds F-actin, suggesting that Spindly serves as a link between dynein and cortical actin in axons. Therefore, Spindly plays a critical role during neurodevelopment by mediating dynein-driven sorting of axonal microtubules.

**Significance Statement:** Neurons send and receive electrical signals through long microtubule-filled neurites called axons and dendrites. One of the main structural differences between axons and dendrites is how their microtubules are organized. Axons contains microtubules with their plus-ends out while microtubules in dendrites are organized with mixed or plus-end-in orientation. Dynein, the main minus-end microtubule motor, anchored to cortical actin filaments in the axons is responsible for the uniform microtubule polarity in axons. However, it is unknown how dynein is recruited to the actin cortex in axons. The major finding of this work is that Spindly, a protein involved in anchoring dynein to kinetochores during cell division, has a second important function in interphase cells recruiting dynein to the actin cortex in axons.

## Introduction

Neurons are post-mitotic cells that transmit electrical signals through long neurites called axons and dendrites. Electric signals captured by dendrites are sent unidirectionally through axons and transmitted to other cells. The polarity of microtubules is strikingly different in these two types of neuronal processes. Axons contain microtubules oriented predominantly with their plus-ends-out, while dendrites contain a large fraction of minus-end-out microtubules (1, 2). Failure in establishing correct polarity of microtubules results in defects of cargo sorting (3). The differences in microtubule orientation between axons and dendrites are gradually established during development. In cultured early stage neurons, non-polarized neurites contain microtubules with mixed orientation. Later the neurite that becomes an axon reorganizes its microtubule network from mixed to uniform polarity with their plus-ends-out (4, 5). These observations suggest that there is a dedicated mechanism for sorting microtubules that selectively eliminates minus-end-out microtubules from the axons, however, the molecular basis of this process is unknown. Uniform microtubule polarity in axons can be disrupted either by inactivation of cytoplasmic dynein (3, 4, 6, 7) or by depolymerization of actin microfilaments. (4). Furthermore, F-actin requirement for microtubule sorting can be bypassed by direct recruitment of cytoplasmic dynein to the plasma membrane (4). These data indicate that cytoplasmic dynein linked to the cortical actin filament network sorts microtubule in axons, however, how dynein is tethered to the cortex in axons is unknown.

Dynein/dynactin is the major minus-end-directed motor complex involved in transport of many different cargos along microtubules [reviewed in (8)]. In addition to organelle transport, dynein/dynactin plays an important role in mitosis, positioning the bipolar spindle and driving chromosome segregation (9–11). To accomplish these different tasks, the dynein/dynactin complex associates with protein adaptors that promote its interaction with specific receptors. One of these adaptors is the kinetochore protein Spindly, which recruits dynein/dynactin to the outer plane of the kinetochore in early prometaphase (12, 13). The stabilization of this complex promotes cell-cycle progression by silencing the spindle assembly checkpoint. In interphase, Spindly can be found proximal to focal adhesions at the leading edge of migrating human cultured cells and its depletion impairs cell migration (14, 15). In agreement with the role of Spindly in cell migration, changes in Spindly levels also affects border cell migration in *Drosophila* ovaries (16).

In this work we report for first time a key role of Spindly in development of *Drosophila* neurons. We demonstrate that post-mitotic depletion of Spindly in *Drosophila* neurons causes large-scale neurodevelopmental defects including disruption of the uniform microtubule orientation in axons, mistargeting of axons, and defects in coordination and locomotion in flies. These phenotypes can be fully rescued by expressing full-length Spindly but not by variants with mutations in the dynein-binding domain. In contrast, the kinetochore binding domain of Spindly is dispensable for its neuronal function. Finally, *in vitro* co-pelleting experiments and microscopy data showed that Spindly interacts with F-actin. Therefore, in addition to the well-established role of Spindly in mitosis, we propose that Spindly has an important role in post-mitotic neurons, targeting dynein to cortical actin in axons. Recruitment of dynein to F-actin allows for dynein-dependent microtubule sorting, thus establishing uniform microtubule polarity in axons and proper axon targeting.

## Results

### Spindly is required for microtubule sorting in processes of cultured S2 cells

In our search for proteins that mediate dynein-dependent sorting of microtubules in cell processes, we first conducted a candidate-based RNA interference (RNAi) screen in *Drosophila* S2 cells. S2 cells treated with 2.5 μM of Cytochalasin D (CytoD), an F-actin severing drug, form long microtubule-filled processes (17). Importantly, processes in CytoD treated cells contained abundant cortical actin (Fig. S1A), as CytoD at this concentration induces severing, but not depolymerization of actin filaments (18, 19). The microtubules in these processes have their plus ends pointing away from the cell body, the same orientation as in axons (4). We have previously demonstrated that inactivation of cytoplasmic dynein in S2 cells, similarly to neurons, causes multiple “minus-end-out” microtubules to appear in processes (4). Furthermore, microtubule-sorting activity of dynein required F-actin, as global depolymerization of F-actin in S2 cells by high concentrations of Latrunculin B also resulted in appearance of minus-end-out microtubules in S2 cell processes (4). We, therefore, performed an initial screen for factors defining microtubule orientation in processes using S2 cells with subsequent validation of positive hits in neurons. Our goal in this screen was to identify adaptors that could play a role in the dynein-dependent pathway that results in uniform microtubule orientation in processes. For this candidate-based screen, the following specific groups of proteins that are expressed in S2 cells were tested (see Table S1 for the complete list): a) proteins enriched in the axon initial segment (AIS), a compartment formed in the proximal section of the axon that separates somato-dendritic and axonal compartments in neurons; b) mitotic adaptors that recruit dynein/dynactin to the cortex during cell division; c) cell adhesion proteins that are enriched at the plasma membrane; d) actin-related proteins that regulate actin dynamics; e) dynein adaptors at the kinetochore. As a marker of microtubule polarity, we used a microtubule minus-end binder CAMSAP3 (20) tagged with mCherry. In control cells mCherry-CAMSAP3 is typically scattered along the length of the processes, (Fig. 1A and 1B); only a small fraction of control cells showed accumulation of CAMSAP3 at the tips of the processes (4). Knockdown of dynein heavy chain (DHC) resulted in dramatic accumulation of CAMSAP3 at the tips of the processes (Fig. 1A and 1B). Among all of the candidates tested in this screen, the dsRNA targeting Spindly was the only treatment that, similarly to DHC RNAi, caused massive accumulation of mCherry-CAMSAP3 at the tips (Fig.1A-C). The effect of Spindly dsRNA treatment on microtubule polarity was further verified by visualization of EB1-GFP comets, a plus-end microtubule marker. In control cells, only 5% of the EB1-comets were directed toward the cell body (Fig. 1D and 1E; Video 1). The percentage of plus-end-in EB1-comets was dramatically increased upon treatment with Spindly dsRNA, further confirming that this treatment caused accumulation of minus-end-out microtubules in S2 cell processes (Fig. 1D and 1E; Video 1). The mixed microtubule orientation found in processes of Spindly RNAi cells was not due to depletion of dynein, as western blotting confirmed that Spindly dsRNA treatment reduced protein levels of Spindly (Fig. S1B and S1C) but not cytoplasmic dynein (Fig. S1B and S1C). Consistent with this observation, transport of peroxisomes that requires cytoplasmic dynein (21) was not affected after Spindly knockdown (Fig. S1D, Video 2).

**Figure 1.**
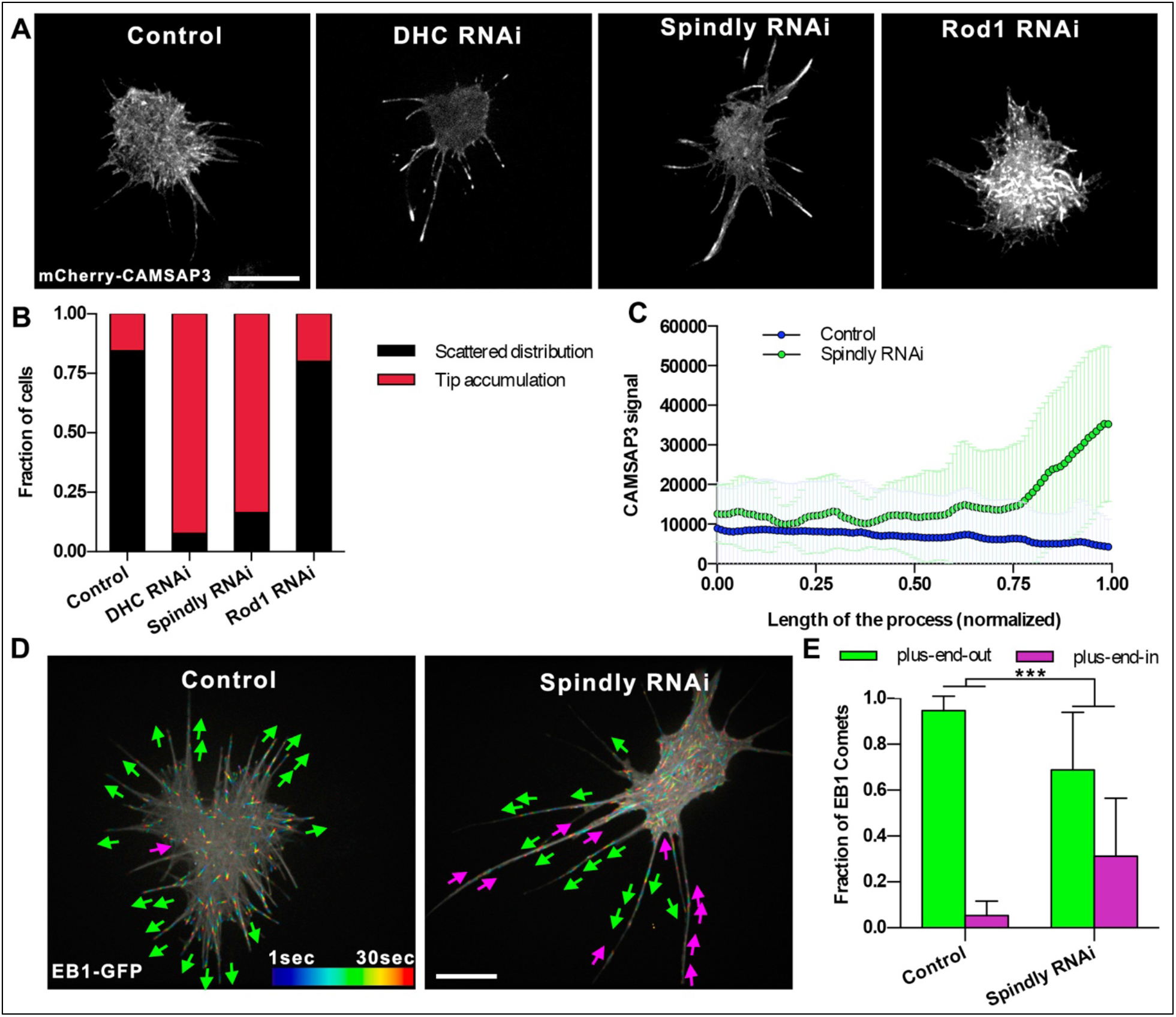
Depletion of Spindly induces mixed polarity in S2 cell processes. A) Representative images of S2 cells expressing mCherry-CAMSAP3 and treated with 2.5μM CytoD to induce the formation of processes. From left to right, control (untreated cells), DHC RNAi, Spindly RNAi and Rod1 RNAi. Note that depletion of DHC or Spindly causes accumulation of CAMSAP3 signal in the tip of the processes. Scale bar, 10 μm. B) Quantification of CAMSAP3 distribution in S2 cells. Cells were classified in two groups; *scattered distribution*-CAMSAP3 signal localizes scattered through the cell; *tip accumulation*-CAMSAP3 signal accumulates in the distal tip of the processes. C) Profile plots of mcherry-CAMSAP3 distribution in S2 cell processes. Fluorescent signal of CAMSAP3 was measured from the proximal (L=0) to the tip (L=1) of the processes in control and Spindly RNAi cells. (Control, n=53 processes; Spindly RNAi, n=62 processes). Data collected from three independent experiments. Error bars represent s.d. D) Representative temporal color code images of control (untreated) or Spindly RNAi S2 cells expressing EB1-GFP. Magenta and green arrows indicate plus-end-in or plus-end-out of EB1 comets, respectively (see Video 1). Scale bars, 10 μm. E) Graphs depicting the direction of EB1-GFP comets in the processes of untreated (control) and Spindly RNAi. (Control, n=20 processes; Spindly RNAi, n=20 processes). Data collected from two independent experiments. Error bars indicate s.d. ***p<0.001

During cell division, Spindly stabilizes dynein/dynactin binding to the kinetochore-binding complex Rod1–Zwilch-ZW10 (RZZ) (see model in Fig. 4J, left panel) (12).To test if the RZZ complex is also involved in regulating microtubule polarity, we treated S2 cells with dsRNA targeting Rod1, but this treatment had no effect on microtubule polarity in S2 cells (Fig. 1A and 1B). Together, these data show that Spindly, but not its closest binding partner in the kinetochore, controls microtubule polarity in S2 cell processes.

### Spindly binds to cortical actin

Next, we tested if Spindly can bind actin microfilaments. To test this interaction, we performed *in vitro* F-actin co-pelleting assays. S2 cell extracts (treated with 2.5 μM CytoD) were incubated in the presence of purified F-actin, followed by high-speed centrifugation to pellet the actin filaments (Fig. 2A). As a negative control, before making extracts, S2 cells were treated with high concentration of Latrunculin B to depolymerize F-actin. To validate this assay, we first demonstrated that in our centrifugation conditions, a-actinin (an actin-binding protein) was enriched in the F-actin pellet. In contrast, Hsc70 (a molecular chaperone) distribution in this assay was independent of the presence or absence of F-actin (Fig. 2B). Similar to a-actinin, Spindly was found in the pellet fraction together with F-actin suggesting an interaction between Spindly and actin (Fig. 2B and C).

**Figure 2.**
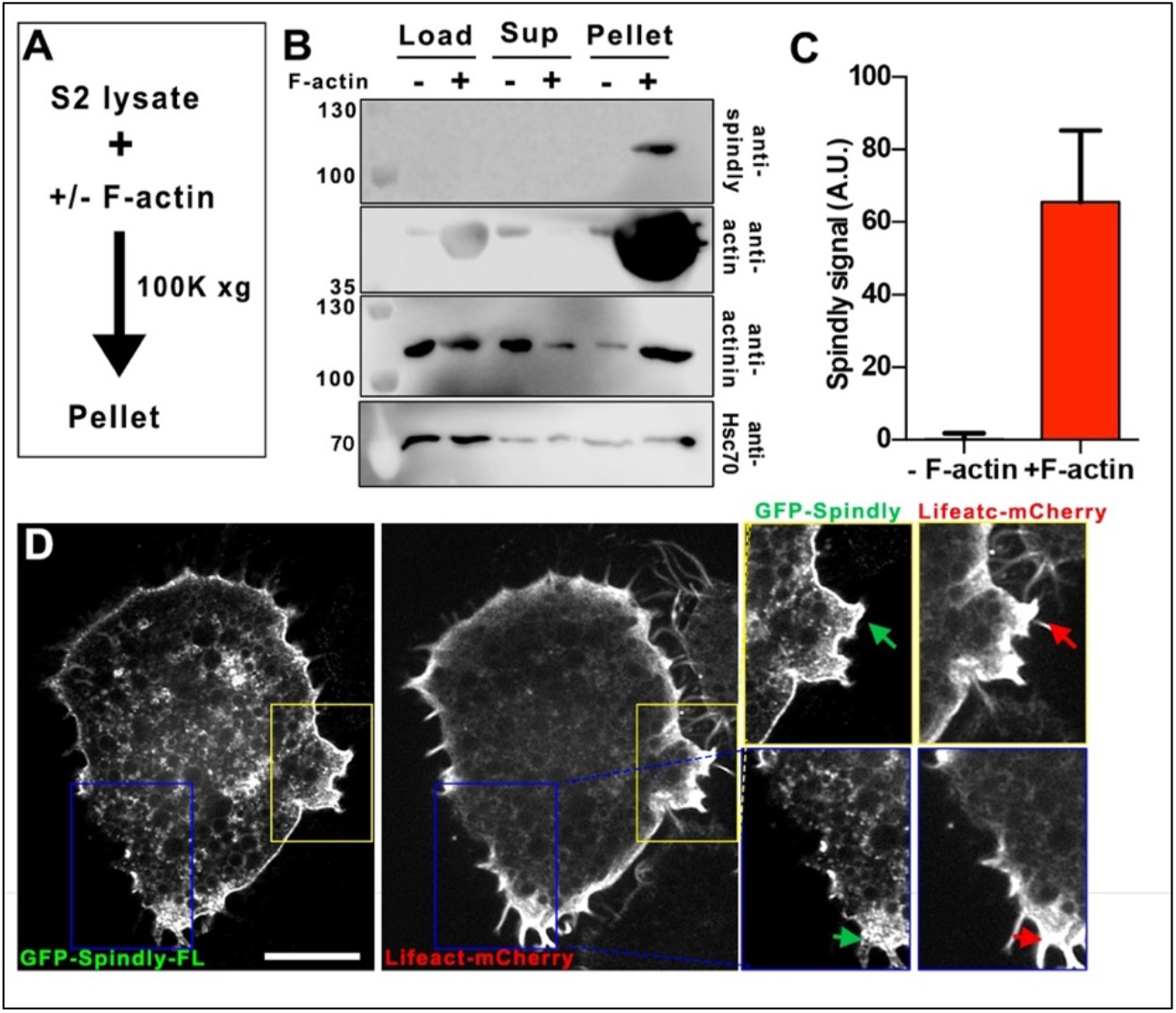
Spindly interacts with F-actin. A) Strategy used for actin co-pelleting experiments. S2 cell lysates were incubated in the presence or absence of F-actin (in the presence of 2.5μM CytoD or 10μM LatB, respectively) and then samples were centrifuged at 100k *xg* for subsequent analysis by western-blot B) Western-blot analysis of Spindly, actin, α-actinin and Hsc70 levels from samples treated as described in Fig 1F. Cell extracts (load), supernatants (sup) and pellets are shown in the membrane. Note that Spindly was only detected in pellets of samples incubated with F-actin. C) Graph depicting the levels of Spindly pulled down in the pellets from experiments shown in Fig. 1G. Data obtained from three independent pull-down experiments. Error bars represent s.d. D) Representative confocal images of a S2 cell expressing GFP-SpindlyFL and Lifeact-mCherry. Note that there is an important accumulation of Spindly in the cortex. The yellow are blue boxes correspond with the areas shown in the right panels. Arrows point to regions where there is a colocalization of Spindly and actin Scale bar, 10 μm.

To further investigate this interaction between Spindly and F-actin, we expressed GFP-SpindlyFL (full-length) together with an actin-marker (Lifeact-mCherry) in S2 cells. Strikingly, a significant fraction of Spindly was highly enriched in the actin-containing cortex (Fig. 2D). These data further support the idea that Spindly interacts with actin filaments and suggest that it could function as a linker between the actin cortex and cytoplasmic dynein as it binds both components.

### Spindly controls microtubule polarity in *Drosophila* axons

Based on our S2 cell data, we suggested that Spindly could regulate uniform microtubule polarity in axons. To test this hypothesis, we depleted Spindly from *Drosophila* neurons using shRNA driven by the post-mitotic pan-neuronal driver elav-Gal4 (22), thus avoiding any effects of Spindly depletion on neuroblast division. We created a transgenic fly stock containing elav-Gal4 and Ubi-EB1-mCherry. By crossing these flies with UASp-Spindly-shRNA, we can visualize EB1-mCherry comets in primary neuronal cultures isolated from third-instar larval brain in the Spindly RNAi background. Quantification of the EB1-comet direction showed that in 48 hour-old cultures, control neurons developed axons with at least 80% of their EB1-comets moving toward the tips of the processes, indicating that most axonal microtubules, as expected, have plus-end-out orientation (Fig. 3A; left panel, Fig. 3C). Depletion of DHC induced the formation of mixed polarity microtubule arrays in axons, as EB1-comets travelled in both directions (53% and 47% plus-end-out and plus-end-in, respectively) (Fig 3A; middle panel, Fig. 3C), consistent with our previously published data (4). Depletion of Spindly, similarly to DHC depletion, caused the EB1 comets to move in both directions (64% and 36% plus-end-out and plus-end-in, respectively) (Fig 3A; right panel, Fig. 3C).

**Figure 3.**
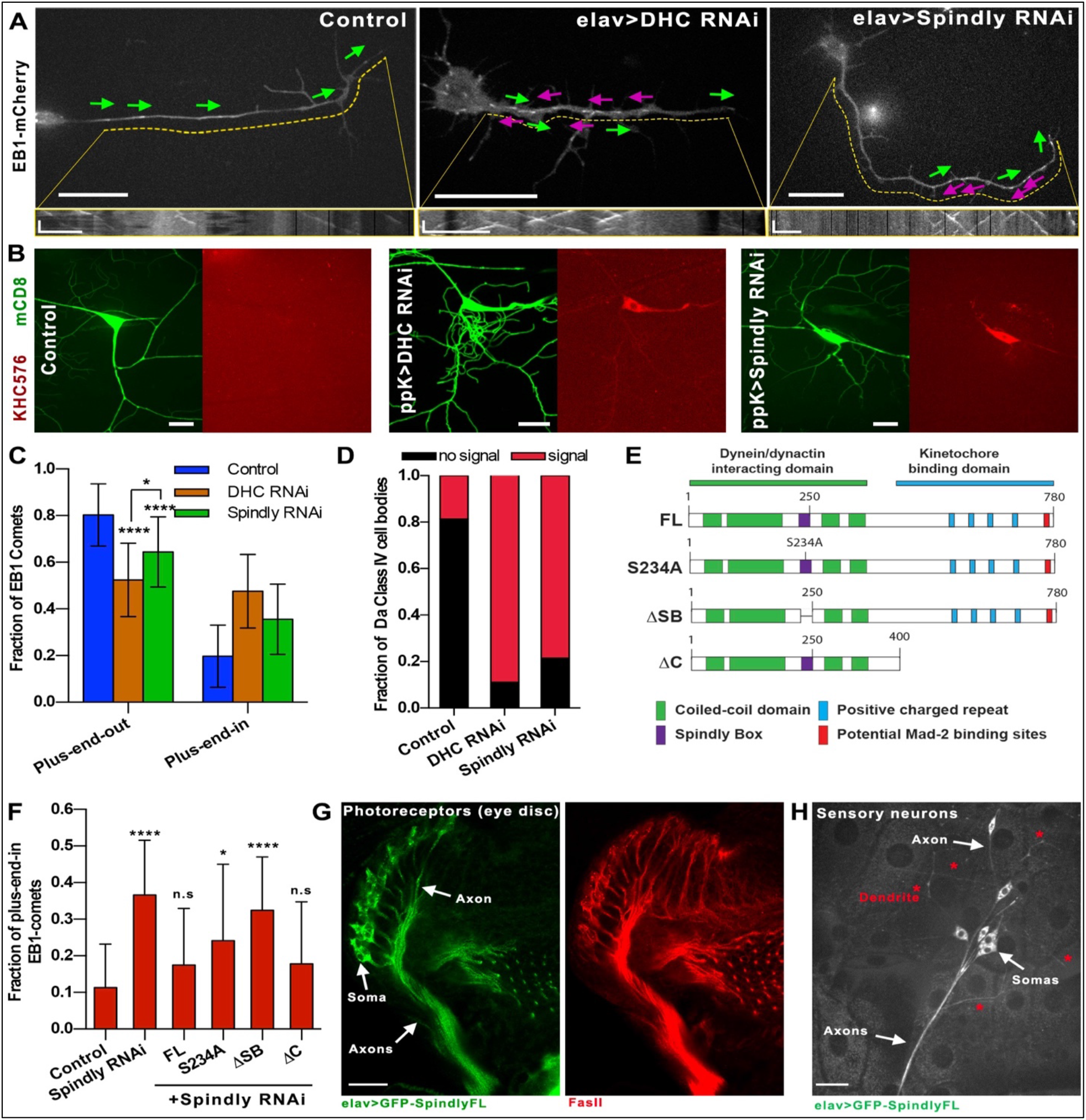
Spindly is required for sorting of microtubules in axons. A) Representative still images of EB1-mCherry expressing primary neurons cultured for 48 hr. Magenta and green arrows indicate the direction of EB1 comets in the axons (plus-end-in and plus-end-out, respectively) (Video 3). Kymographs of EB1 comets are shown below corresponding images. Dashed yellow lines define the area of the axon used for plotting EB1-GFP kymographs. Horizontal scale bars, 10 μm in the main panels and 5μm in the kymographs, respectively. Vertical bars in the kymographs represent time (60 sec). B) Representative Z-stack images of class IV sensory neurons of control (ppk-Gal4; left panels) or ppk>DHC RNAi (middle panels) or ppk>Spindly RNAi (right panels). Membrane and microtubule plus-ends were labeled with mCD8-GFP and KHC(1-576)-RFP, respectively. Scale bar, 20 μm. C) Graph depicting the fraction of axonal EB1-GFP comets directed toward the tip of the process (plus-end-out) or the cell body (plus-end-in). Data obtained from three individual experiments (control, n=33 axons; DHC RNAi, n=26 axons; Spindly RNAi, n=31 axons). Error bars represent s.d. *p=0.0134; ****p<0.0001 D) Classification of sensory neurons based on the KHC(1-576)-RFP signal intensity in the somas (control, n=29 sensory neurons; DHC RNAi, n=16 sensory neurons; Spindly RNAi, n=20 sensory neurons). E) Schematic cartoon showing the full-length *Drosophila* Spindly protein and variants used for Spindly RNAi rescue assays in flies. Note that all these Spindly proteins are resistant to the shRNA sequence used to knockdown Spindly [Figure adapted from (16)]. F) Fraction of microtubules with plus-end-in found in Spindly RNAi axons from neurons expressing full-length Spindly and variants (S234A, ΔSB and ΔC). Data obtained from at least three independent experiments (number of quantified axons for each genotype ranged between 20 and 74). n.s. non-significative, *p=0.011, ****p<0.0001. G) Confocal image of a fixed third-instar eye disc expressing elav>GFP-SpindlyFL (left panel) and immunostained with Fas-II (right panel). Spindly is enriched in both somas and axons of the photoreceptor neurons. Scale bar, 10 μm. H) Sensory neurons expressing elav>GFP-SpindlyFL. Notice that Spindly localizes primary in the somas and axons. A small fraction of Spindly can be found in dendrites. Scale bar, 20 μm.

Next, we examined the effects of Spindly knockdown on microtubule polarity in neurons *in vivo* using class IV dendritic arborization (da) sensory neurons in third-instar larvae. These neurons are located at the body wall of the animal, making them easy to image *in vivo* (see Fig. S2A). To visualize microtubule polarity in these neurons, we used the RFP-tagged constitutively active motor dimer of kinesin-1, KHC(1-576) (the *Drosophila* version of mammalian KIF5B(1-560)), which accumulates at the plus end of microtubules (23). We used a membrane marker (mCD8-GFP) to identify class IV da neurons. Expression of both markers in flies was driven by a specific class IV neuron driver (ppk-Gal4).

In control class IV da neurons, KHC(1-576)-RFP accumulated at the tips of axons at the ventral nerve cord (data not shown) but was essentially absent from the soma and dendrites (Fig. 3B; left panel, Fig. 3D). Depletion of DHC caused a strong accumulation of KHC(1-576)-RFP in the cell body in 91% of the examined class IV da neurons (Fig. 3B; middle panel). Depletion of Spindly mirrored the KHC(1-576)-RFP accumulation in the soma found in DHC RNAi class IV neurons (78% of the examined neurons),demonstrating that there is a significant fraction of axonal microtubules oriented with their plus ends facing the cell bodies (Fig. 3B; right panel, Fig. 3D). Together, these data demonstrate that Spindly is required for uniform orientation of axonal microtubulesin culture and *in vivo*.

### Dynein-binding, but not kinetochore-binding domain of Spindly is required for sorting axonal microtubules

We next investigated which of Spindly’s domains was required for its function in microtubule shorting in axons. Spindly contains two main functional domains (Fig. 3E). Its amino-terminal domain is a rod-like structure that includes the binding motifs for dynein light intermediate chain and the pointed-end complex of dynactin (24). One of these motifs, the highly conserved Spindly Box, is required for Spindly’s ability to recruit dynein/dynactin to the kinetochore (25, 26). In contrast, its carboxy-terminal domain contains four positively charged repeats responsible for the interaction with the kinetochore RZZ complex (Fig. 3E). Disruption of either of these two domains results in failure of cell cycle progression in human cells (26). To identify which domain of Spindly is required for neurodevelopment, we used transgenic flies expressing full-length Spindly and a set of Spindly deletion mutants, all resistant to the Spindly shRNA. Two mutants, Spindly-S234A and Spindly-ΔSB, had disruptions in the dynein-binding motifs (a single amino-acid replacement in the Spindly Box, or the deletion of the entire Spindly Box, respectively) (Fig. 3E). The third mutant, Spindly-ΔC, contained intact dynein/dynactin binding domain but lacked the domain associated with its kinetochore binding in *Drosophila* (Fig. 3E) (16).

To study if the expression of these Spindly variants was capable of rescuing the microtubule polarity defects found in Spindly RNAi axons, we created a fly containing Ubi-EB1-mCherry and UASp-Spindly RNAi, and crossed it with flies expressing elav-Gal4 and UASp-GFP-Spindly (full-length (FL) and variants). Microtubule polarity was examined in axons of cultured neurons obtained from third-instar larva brains of these flies (as in Fig. 3A). The microtubule polarity defects observed in axons of Spindly RNAi neurons was fully rescued by ectopic expression of GFP-tagged Spindly-FL, further confirming that microtubule polarity defects observed after Spindly knock-down were not caused by an off-target effect of the shRNA (Fig. 3F). Expression of Spindly-S234A or -ΔSB, deficient in dynein interactions, failed to rescue the Spindly knock-down phenotype (Fig. 3F). In contrast, expression of Spindly-ΔC, the variant that does not bind kinetochores, resulted in generation of axons with uniform microtubule polarity. These data indicate that dynein-binding, but not kinetochore-binding activity of Spindly is required for microtubule organization in axons (Fig. 3F).

### Spindly accumulates mainly in the soma and axons of neurons

To better understand the function of Spindly in neurons, we examined the distribution of GFP-SpindlyFL using neuron-specific drivers. This fusion protein was fully active as it rescued the microtubule polarity defects found in axons of Spindly RNAi neurons (Fig. 3F). We first examined the photoreceptor neurons in the eye disc of dissected third-instar larva brains. Photoreceptor neurons extend bundles of axons converging into a classic umbrella-like (retinotopic pattern) to the optic stalk (see Fig. 4A). To visualize these structures, dissected brains form third-instar larvae expressing elav>GFP-SpindlyFL were fixed and stained with a Fasciclin II (Fas-II) antibody that specifically labels axons. We found that in these neurons GFP-Spindly was enriched both in cell bodies and axons (Fig. 3G). The same distribution was observed using the photoreceptor-specific driver ninaE-GRM-gal4 (data not shown). We also examined the distribution pattern of GFP-Spindly in da sensory neurons *in vivo*. As in the case of photoreceptor neurons, GFP-SpindlyFL localized mainly in cell bodies and axons, with only weak GFP-Spindly signal seen in sensory neuron dendrites (Fig 3H). These data support that Spindly expression in neurons is preferentially enriched in cell body and axon.

**Figure 4.**
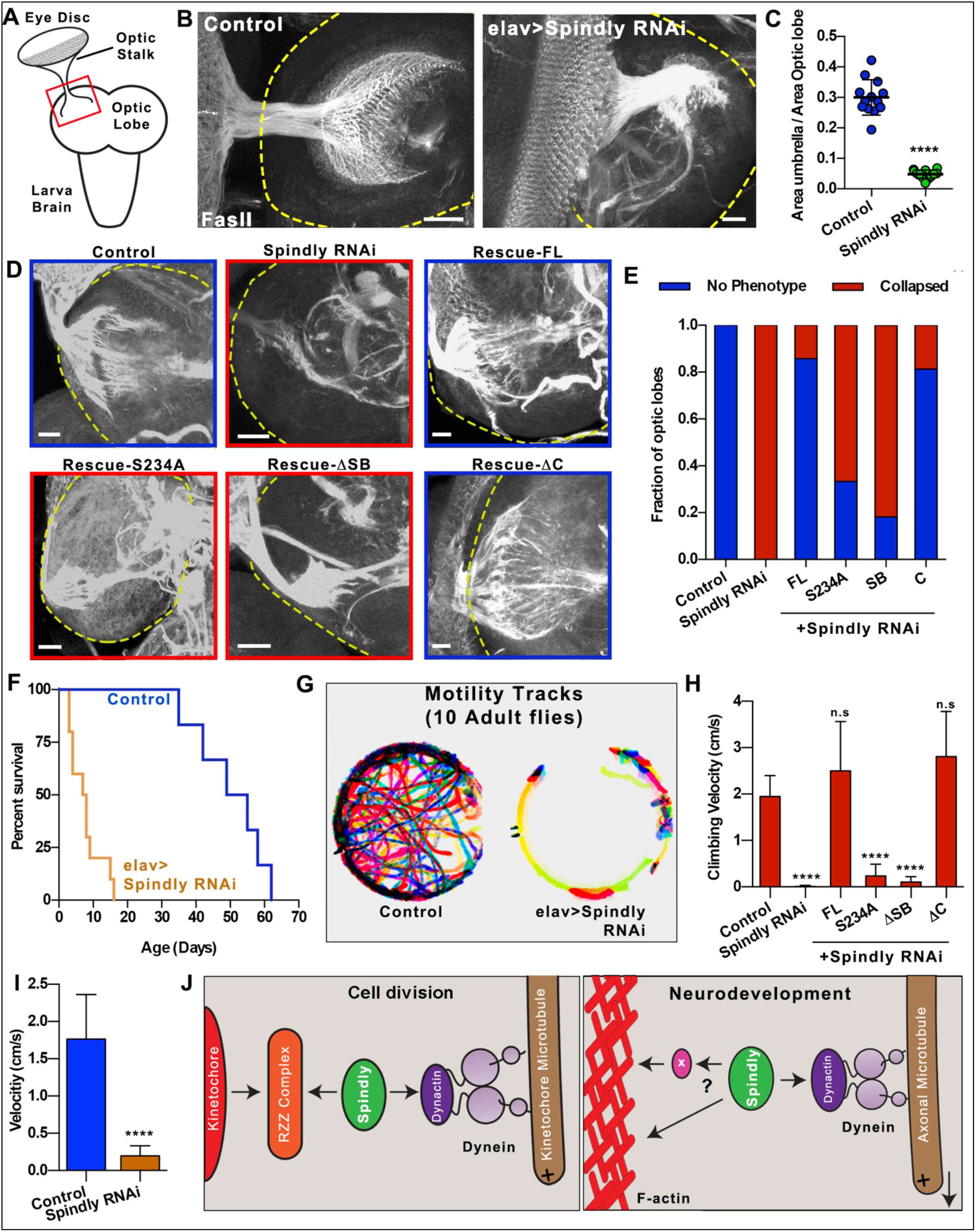
Depletion of Spindly causes mayor defects in neurodevelopment. A) Diagram of dissected third-instar larval brain. The optic stalk, which contains axons of photoreceptors neurons, connects the eye imaginal disc to the optic lobe of the brain. B) Third-instar larval brains were fixed and stained with anti-fasII antibody. Axon terminals at the optic lobe display a well-spread umbrella-like pattern in control larvae (left panel). This pattern is collapsed in all Spindly RNAi larvae (right panel). Yellow dashed-lines highlight the boundary of the optic lobes. Scale bar, 20 μm. C) Quantification of the area of the optic lobe innervated by the optic stalk. (Control, n=13 brains, elav>Spindly RNAi, n=12 brains) ****p<0.0001 D) Representative examples of third-instar dissected brains fixed and immunostained with Fas-II from control, elav>Spindly RNAi and Spindly replacement genotypes (elav>Spindly RNAi + GFP-Spindly variants). Yellow dashed-lines highlight the boundary of the optic lobes. Scale bars, 20 μm. E) Quantification of the retinotopic innervation in the optic lobes from experiment shown in (D). Innervation pattern was classified in two groups: “no phenotype” when innervation was fully extended as observed in control brains; “collapsed” when photoreceptors failed to properly innervate the optic lobe as observed in Spindly RNAi brains. Data obtained from at least 10 optic lobes per genotype. F) Lifespan of control (elav-Gal4) and elav>Spindly RNAi adult flies. Data obtained from 10 individual flies per each genotype. G) Temporal color-coded tracks of 10 flies per condition for 100 sec (see Video 4). H) Quantification of climbing assays. Climbing velocities of control and Spindly RNAi flies coexpressing the full-length Spindly and mutants (see Video 5). n.s. non-significative; ****p<0.0001 I) Graph depicting the locomotion velocity of control (elav-Gal4) and elav>Spindly RNAi flies measured from Video 4 (10 flies for control and elav>Spindly RNAi). ****p<0.0001 J) Model of dynein recruitment to the cortex mediated by Spindly. Left panel. Established role of Spindly during cell cycle progression. Spindly is responsible for stabilizing the interaction of dynein/dynactin complex to the out-layer of the kinetochore. The stabilization of this complex promotes cell-cycle progression by silencing the spindle assembly checkpoint [figure modified from (12)]. Right panel. Novel function of Spindly role during neurodevelopment presented in this work. Spindly is responsible for recruiting of dynein/dynactin complexes to filaments of actin (F-actin) in the axon. Spindly can recruit dynein to F-actin through a direct interaction to F-actin or through intermediate adaptors. Recruitment of dynein/dynactin to F-actin activates microtubule transport powered by dynein. Note that the direction of the moving microtubule is dictated by the microtubule orientation. If a microtubule has a plus-end-in orientation, this microtubule will be transport towards the soma resulting in microtubule polarity sorting. In contrast, if the microtubule has a plus-end-out orientation the microtubule will be transport towards the tip of the axon promoting axon outgrowth.

### Depletion of Spindly affects axonal, but not dendritic patterning

We next studied if disruption of microtubule polarity observed in axons of Spindly RNAi neurons affects the development of the *Drosophila* nervous system. We inspected the photoreceptor axon-targeting pattern in the optic lobes of the visual system of third-instar larvae. The area of the optic lobe covered by the photoreceptor axons was used to quantify the integrity of the retinotopic pattern (Fig. 4A). In a control brain, the axons from photoreceptor neurons projected a retinotopic pattern that covered 30% of the optic lobe (Fig. 4B and 4C). Post-mitotic depletion of Spindly in neurons consistently caused a dramatic collapse of these structures (Fig. 4B; right panel). In brain from Spindly RNAi flies, the photoreceptor stalk covered only 4.7% of the optic lobe (Fig. 4C). These data demonstrated that proper axon development was grossly impaired in the absence of Spindly. In good agreement with the microtubule polarity rescue assay, the expression of the Spindly-FL and Spindly-ΔC, but not Spindly-S234A and Spindly-ΔSB, fully rescued the retinotopic collapse in the optic lobe of brains of third-instar larval induced by Spindly RNAi (Fig 4D and 4E).

Dynein is not only important for axon development, but its activity is also required for dendritic arborization in Class IV sensory neurons. DHC depletion caused major defects in dendrite development (Fig. S2A and S2B) (3). Both Sholl analysis and dendrite length quantification showed that dynein depletion affects proper dendritic arborization (Fig. S2C and S2D) (3, 7). We asked if Spindly, in addition to its role in proper axon development, is also required for dynein function in dendrite development. Interestingly, elav>Spindly RNAi third-instar larvae did not show any defects on dendritic arborization development, as their primary and secondary dendritic branches were fully developed (Fig. S2B-D). Together, these data further support the idea that Spindly function (as its localization) is primarily restricted to axons.

### Depletion of Spindly compromises locomotion and survival rates in adult flies

Despite axonal microtubule polarity and pattering caused by Spindly knock-down, these larvae developed to adulthood. However, elav>Spindly RNAi flies eclosed from the pupal case displayed severe locomotion and coordination defects as well as a shortened lifespan (Fig. 4F and 4G, and Video 4). Locomotion velocity of Spindly RNAi flies were significantly lower than control flies (1.7 cm/s and 0.2 cm/s, control and elav>Spindly RNAi, respectively) (Fig. 4I and Video 4). The climbing defects found in Spindly RNAi flies were fully rescued by Spindly-FL and Spindly-ΔC, but not by the Spindly variants with mutations in the Spindly Box domain (Fig 4H and Video 5). Together, these data indicate that Spindly plays important post-mitotic roles in neurodevelopment, and these activities require its interaction with dynein, while its kinetochore-binding activity is dispensable for these functions.

## Discussion

Several works using different model systems have demonstrated that uniform polarity of microtubules in axons requires activity of cytoplasmic dynein (3, 4, 6, 7) recruited to cortical actin filaments (4). However, the mechanism that targets dynein to cortical actin remained unknown. In the search for adaptors involved in the recruitment of dynein to F-actin, we performed a targeted RNAi screen and showed that Spindly, a well-characterized protein that recruits dynein to kinetochores in mitosis, is required in post-mitotic neurons for dynein-dependent organization of microtubules in axons. Depletion of Spindly in *Drosophila* neurons impairs axonal microtubule sorting; brain of Spindly-depleted third-instar larvae showed severe defects in axonal patterning. These neurodevelopmental defects result in impairment of coordination and locomotion, and reduced lifespan of adult flies. These phenotypes are not caused by reduction of the dynein level or inhibition of dynein-driven organelle transport upon Spindly knockdown. Spindly RNAi defects found in the *Drosophila* brain are fully rescued by expression of full-length Spindly or the variant deficient in kinetochore binding, but not by variants with mutations in its dynein binding domain. Together, these data suggest that Spindly plays a role in neurodevelopment through a dynein-dependent pathway.

Spindly was originally identified as a mitotic component recruited to the kinetochore in a RZZ-dependent pathway (12). In mitosis, the formation of stable interactions between kinetochores and dynein in the metaphase plate are required to silence spindle assembly checkpoint, allowing the progression of the cell cycle to anaphase (Fig.4J; left panel) (12). In Spindly-depleted cells, dynein motors fail to be recruited to the outer plate of the kinetochores, and the lack of stable kinetochore-microtubule contacts results in cell cycle arrest in metaphase (12, 13, 27). We propose that in post-mitotic neurons Spindly is also important for dynein recruitment. However, in the case of neurons, Spindly recruits dynein to the actin cortex in axons (Fig. 4J; right panel). Importantly, two types of experiments show that this post-mitotic role of Spindly is independent of its canonical role in mitosis. First, depletion of Rod1, one of the kinetochore components that interacts with Spindly during cell division, did not affect the microtubule polarity. More directly, expression of the Spindly variant lacking its kinetochore binding domain (Spindly-ΔC) rescues the Spindly RNAi defects in *Drosophila* neurons. Both observations together suggest that the neuronal spindly-dependent pathway of dynein recruitment and microtubule organization is different from its canonical mitotic pathway.

It has been shown recently that, in addition to its role in cell division, Spindly functions in interphase cells. Both in mammalian cells and in *Drosophila*, changes of Spindly level negatively impact cell migration (14, 16). Remarkably, both Spindly and dynein/dynactin complex are found at the cell cortex at the leading edge of migrating human cells (14). In good agreement with this observation, our biochemical and cellular assays showed that Spindly interacts with F-actin. However, at this point it is unknown whether Spindly interacts with actin directly or whether this interaction is mediated by other proteins. The lack of known/predicted actin-binding domains in Spindly favors the second scenario. It will be very interesting to identify the proteins that form a complex with Spindly in interphase and find components of this complex that are involved in the recruitment Spindly and dynein to F-actin.

Importantly, the loss of dynein activity in *Drosophila* sensory neurons did not affect microtubule polarity in dendrites, indicating that the microtubule-sorting activity of dynein is restricted to axons (7). The apparent restriction of the dynein-recruiting Spindly activity to axons is yet to be determined. Our data support that Spindly primary localizes in the cell body and axon, although a small fraction of the protein can be found in dendrites. It has been reported that Spindly is post-translationally modified and modifications affect its localization and/or functions. For example, farnesylation of Cys602 of human Spindly is essential for its accumulation at prometaphase kinetochores (28, 29). Spindly can also be phosphorylated by CDKs during cell division, and S515 of human Spindly is the major phosphorylation site. Interestingly, this modification seems to regulate ZW10 function rather than Spindly localization (26). We hypothesize that post-translational modifications in the amino-terminal domain of Spindly may regulate its role in neurodevelopment. Recently it has been reported that other kinetochore proteins are important for neurodevelopment (30, 31) [reviewed in (32)]. For example, depletion of Mis12, Knl1 and Ndc80 (other kinetochore components) results in abnormal neuromuscular junctions and central nervous system development both in *Drosophila* and in *C. elegans* (30, 31). However, the precise roles of these kinetochore proteins in neurodevelopment are as yet unknown.

Is cortical dynein the only factor that sort microtubule polarity in axons? Data from a number of groups using different model systems support the idea of dynein being the universal motor that sorts axonal microtubules. However, depletion of other proteins also results in microtubule polarity defects in axons. For example, it has been reported that TRIM46, a microtubule-associated protein anchored to the AIS through AnkG, is able to organize uniformly-oriented microtubule bundles (33). Obviously, our S2 screen is not comprehensive, and we cannot even exclude that the targets that gave us negative results in S2 screen could in fact be involved in microtubule organization in neurons as S2 cell processes are a very crude model of microtubule organization in neurons. It is likely that the development of a non-polarized neurite to a fully functional axon is a complex process that requires cooperation of multiple components including dynein, dynein adaptors, other microtubule binding proteins and components of the AIS. Our data shown here simply demonstrate that Spindly belongs to this group of proteins and is an important factor that in the recruitment of dynein to F-actin. Future work will show how these “mitotic” components work together to properly organize axonal microtubules.

## Acknowledgements

We thank Dr. Anna Kashina, Dr. Jill Wildonger, Dr. Masha Gelfand and Dr. Rosalind Norkett for critical reading of this manuscript, Dr. Christian Suarez for the purified actin, Dr. H. Okhura for the Ubi-EB1-mCherry fly stock and the Bloomington Drosophila Stock Center (supported by National Institutes of Health grant P40OD018537) for fly stocks. We also thank the members of the Gelfand laboratory for their suggestions and support. Research reported in this study was supported by the National Institute of General Medical Sciences grants R01GM052111 and R35GM131752 to V.I.G.

## Author Contributions

U.dC designed and performed experiments, analyzed data and wrote the paper; H.J.M. designed experiments and drafted the paper; V.I.G designed experiments and wrote the paper

## Material and Methods

### Flies stocks and plasmids

Fly stocks and crosses were cultured on standard cornmeal food based on Bloomington Stock Center’s recipe at room temperature. The following fly stock lines were used in this study: Spindly-RNAi TRiP line (Valium 20, Bloomington stock #34933, 3rd chromosome attP2 insertion, targeting Spindly CDS 1615-1635); DHC64C-RNAi TRiP lines (Valium 20, Bloomington stock #36698, 3rd chromosome attP2 insertion, targeting DHC64C CDS 1302–1322; Valium 22, Bloomington stock #36583, 2^nd^ chromosome attP40 insertion, targeting DHC64C CDS 10044–10064); elav-Gal4 (3^rd^ chromosome insertion, Bloomington stock #8760; 2^nd^ chromosome insertion, Bloomington stock #8765) (34); ppk-Gal4 (2^nd^ chromosome insertion, Bloomington stock #32078); UASp-KHC576-TagRFP (35); UASp-GFP-Spindly constructs (Full-length, S234A, ΔSB and ΔC) (3^rd^ chromosome insertions) (16); w,ubi-EB1-mCherry (X chromosome insertion) (36); yw;ppk-tdtomato;elav-Gal4 (3^rd^ chromosome). For Spindly rescue assays, we created flies expressing yw;elav-Gal4;UASp-GFP-Spindly variants and w,ubi-EB1-mCherry; Spindly-RNAi TRiP were generated using standard balancing procedures.

To study microtubule polarity in *Drosophila* S2 cells, pMT-mCherry-CAMSAP3 and pMT-EB1:EB1-GFP plasmids were used to visualize microtubule minus-ends and plus-ends, respectively (4). To visualize peroxisome movement, S2 cells were transfected with pAC-GFP-SKL (17).

### *Drosophila* cell culture: primary neurons and S2 cells

Primary neurons were obtained from dissected brains of 3rd instar larva as previously described (37). Neurons were plated onto Concanavalin A-coated coverslips in supplemented Schneider’s medium (20% fetal bovine serum, 5 μg/ml insulin, 100 μg/ml penicillin, 100 μg/ml streptomycin, and 10 μg/ml tetracycline). *Drosophila* S2 cells were cultured as previously described (17). S2 cultures were induced to form microtubule-based processes adding 2.5 μM CytoD in the medium. For knockdown assays in S2 cells, cultures at 1.5 x 10^6^ cells/mL were treated twice with 20 μg of dsRNA (day 1 and day 3) and cell analysis was performed on day 5. Double-stranded RNA was transcribed in vitro with T7 polymerase, and purified using LiCl extraction. Primers used to create T7 templates from fly genomic DNA are described in Table 1.

### Immunostaining and antibodies

Optical lobes, dissected from third-instar larva in 1× PBS, were fixed in 4% (wt/wt) formaldehyde (methanol-free) diluted in PBT (1× PBS, 0.1% Triton X-100) for 20 min; optical lobes were then washed five times with PBTB (1× PBS, 0.1% Triton X-100, 0.2% BSA) for 10 min and blocked in 5% (vol/vol) normal goat serum for 1 h. Samples were then incubated with primary anti-Fasciclin II antibody (1:40 of the concentrate antibody, 1D4; Developmental Studies Hybridoma Bank) at 4C overnight. Samples were then washed five times with PBTB (10min each wash) and incubated with secondary anti-mouse for 2h at RT. Finally, samples were washed with PBT 10 min five times before mounting.

For western-blots, the following antibodies were used: anti-Spindly antibody (12); anti-actin JLA20 (deposited to the DSHB by Lin J.J) (38); anti-alpha-actinin 2G3-3D7 (deposited to the DSHB by Saide, J.D) (39); anti-Hsc70 (K-19; Santa Cruz Biotechnology) and anti-DHC monoclonal antibody 2C11-2 (40).

### Microscopy and image acquisition

To image EB1-GFP, EB1-mCherry, mCherry-CAMSAP3, KHC(1-576)-RFP, GFP-SKL in *Drosophila* primary neurons, da sensory neurons and S2 cells a Nikon Eclipse U2000 inverted microscope equipped with a Yokogawa CSU10 spinning disk head, Perfect Focus system (Nikon) was used with a 100 × 1.45 or 40x 1.30 oil immersion lenses. Images were acquired using Evolve EMCCD (Photometrics) and controlled by Nikon Elements 4.00.07 software. For EB1 and peroxisome transport time-lapses, images were collected every 2sec for either 1 or 2min. Immuno-stained brains were imaged using a Nikon Ti2 inverted microscope equipped with a A1plus scanning confocal and a GaAsP detector using a 40x 1.30 NA oil lens. For GFP-Spindly localization assays, samples were imaged in a Nikon Ti2 inverted microscope equipped with a W1 confocal spinding disk, a Live-SR (Gataca systems) and a EMCCD Prime 95B (Photometrics) sensor.

### Assays with adult flies

For survival analysis, newly eclosed flies were collected and housed at a density of 20 flies per vial in apple-juice agar supplemented with dry yeast. Flies were flipped to a fresh vial every 3 days. For motility and climbing assays, newly eclosed flies were transfer into a 35mm dish (motility) or vial (climbing). 1h after collection, the motility of the flies was recorder with a DSLR camera.

### Image Quantification

mCherry-CAMSAP3 distribution in S2 cell processes was quantified using plot-profiles. The distance from the proximal to distal tip in the plot profile was normalized using a custom MATLAB program. Quantification of directions of EB1 comet in axons of primary neurons and in S2 cell processes was performed using a temporal-code plugin in FIJI. Peroxisome transport and adult fly motility was quantified using the automated particle-tracking software (41). Z-stack images of Class IV sensory neurons expressing ppk-tdTomato were stacked (maximum intensity projection) and then masked with Curve Tracing (Carsten Sterger’s algorithm) plugin in FIJI. Masked files were then threshold to binary images, and dendritic branching and total dendrite length were quantified using Sholl analysis (42) and pixel counting, respectively.

### Actin co-pelleting experiments

G-actin was polymerized to F-actin as previously described (43). Briefly, purified G-actin (10μM) was incubated with KMEI buffer (50 mM KCl, 1 mM MgCl2, 1 mM EGTA and 10 mM Imidazole (pH 7.0)) for 2h at RT. F-actin was incubated with clarified S2 cell lysates in BRB80 buffer (Na-PIPES 80 mM, EGTA 1mM, MgCl2 1 mM, DTT 1mM (pH 7.0)) for 30min at RT (incubated either with 2.5 μM CytoD or 10 μM LAtB). F-actin and actin-binding proteins were pelleting by ultracentrifugation (100,000 x*g*) at RT. A fraction of the pellets and supernatants were analyzed by SDS-PAGE and western-blot.

### Statistical Analysis

Statistical significance between 2 groups was determined using the unpaired, nonparametric Mann-Whitney test. Data analyses were performed with Prism v6 (GraphPad Software, Inc.).

---

**Figure S1.**
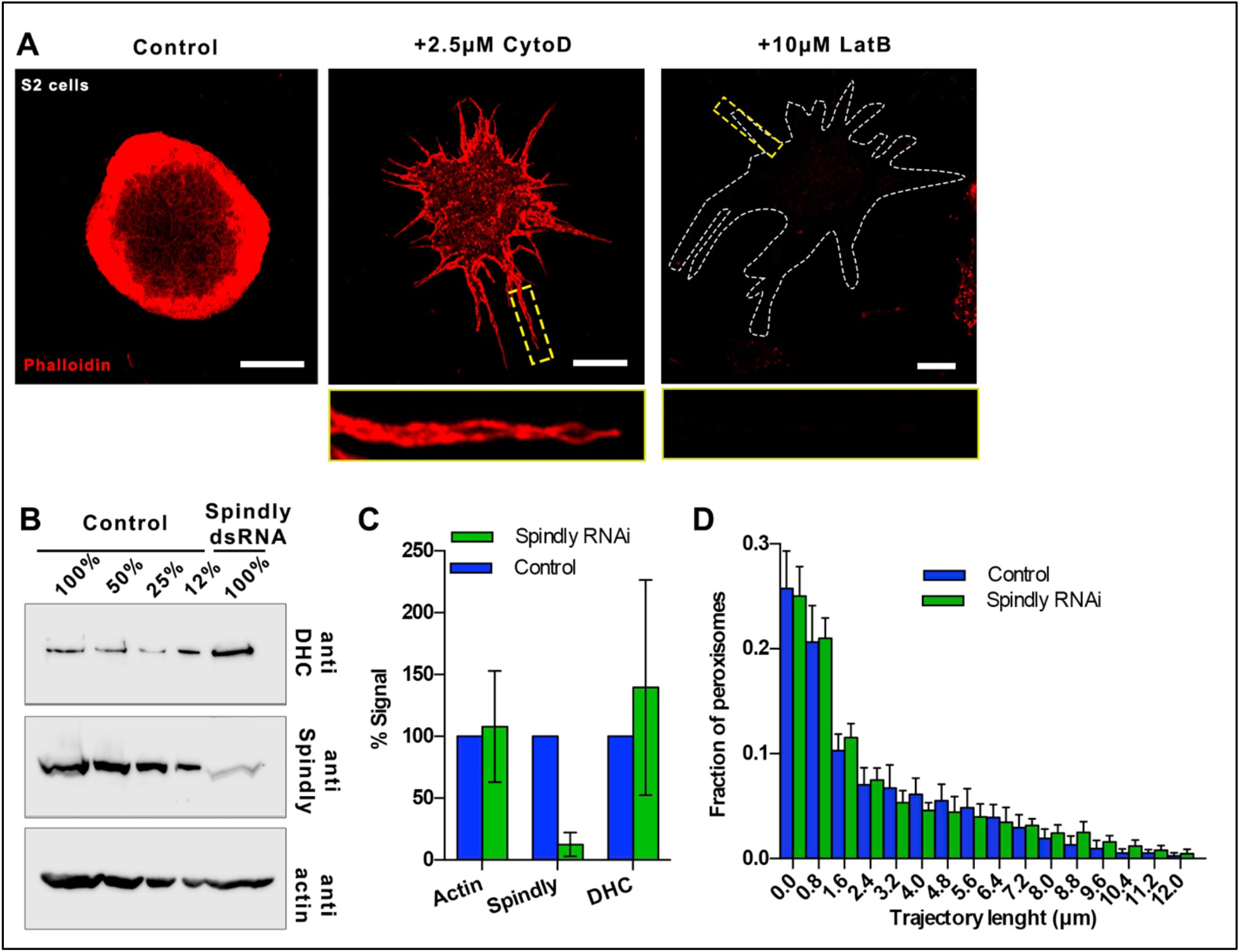
Spindly knockdown does not alter DHC levels. A) Low concentration of CytoD (2.5μM) does not depolymerize cortical actin of S2 processes. S2 cells were plated in the absence of drugs (control) or 2.5μM CytoD or 10μM LatB. F-actin was stained with Rhodamine-Phalloidin. The concentration of CytoD used in the S2 candidate-based screen (2.5 μM) allowed the formation of microtubule-base processes containing cortical actin. In contrast, treatment with high concentration of LatB induced actin depolymerization. Images were taken using the same microscope settings. Scale bars, 10 μm. B) Representative western blots of the S2 lysates of untreated (control) and treated with Spindly shRNA. Dilutions of untreated lysates were provided to estimate degree of knockdown. C) Protein levels were quantified using the western-blots in (B). Data obtained from three independent assays. D) Trajectory length of peroxisome transport in S2 cells untreated (control) and Spindly RNAi (see Video 2).

**Figure S2.**
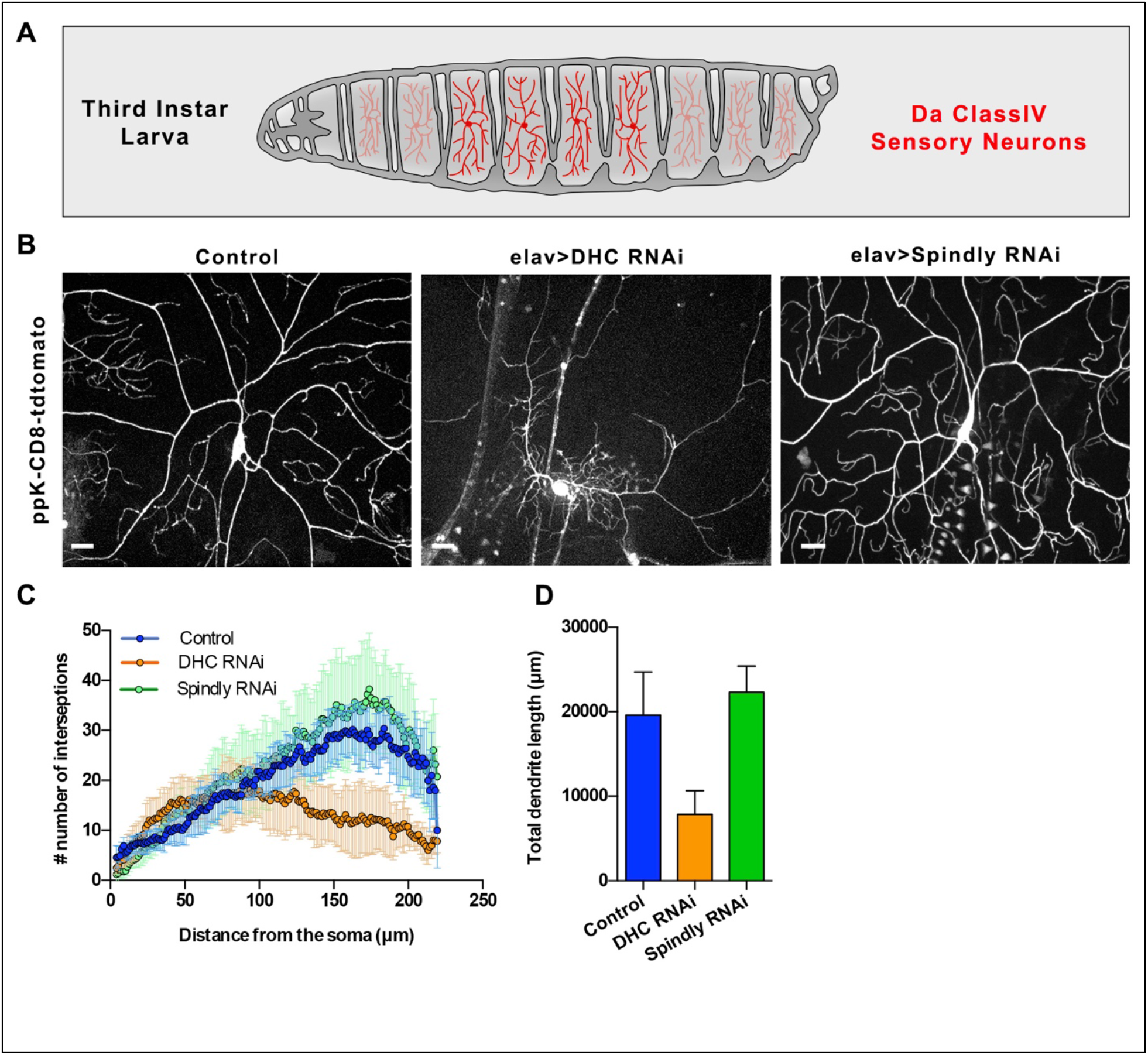
Spindly does not affect development of dendrites. A) Diagram of third-instar larva showing the class IV Da sensory neurons in red. B) Representative max-projection images showing DA neurons labelled with ppk::tdTomato in 3rd instar larvae under control conditions (left panel), DHC RNAi (middle panel) or Spindly RNAi (Left panel). Scale bars, 20 μm. C) Sholl analysis of dendritic arborization of class IV sensory neurons. Data obtained from 12, 13 and 10 animals for control, DHC RNAi and Spindly RNAi, respectively. D) Quantification of the total dendritic length of Da neurons from data obtained in (C).

## Video legends

**Video 1**. Time-lapse of control (untreated cell) or Spindly RNAi S2 cells expressing EB1-GFP. Scale bar, 10 μm. Related to Fig. 1D.

**Video 2.** Time-lapse of control (untreated cell) or Spindly RNAi S2 cells expressing SKL-GFP. Scale bar, 10 μm. Related to Fig. S1D.

**Video 3.** Time-lapse of primary neurons expressing Ubi-EB1-mCherry of three different genotypes. Control (elav-Gal4), DHC RNAi (elav>DHC shRNA) and Spindly RNAi (elav>Spindly shRNA). Scale bar, 10 μm. Related to Fig. 3A.

**Video 4.** Motility assay of control (elav-Gal4) and elav>Spindly RNAi adult flies in a 35 mm dish. Related to Fig. 4G and 4I.

**Video 5.** Climbing assay of adult flies with different genotypes. From left to right: Control (elav-Gal4), Spindly RNAi (elav>Spindly RNAi). Related to Fig. 4H.

**Table S1.**
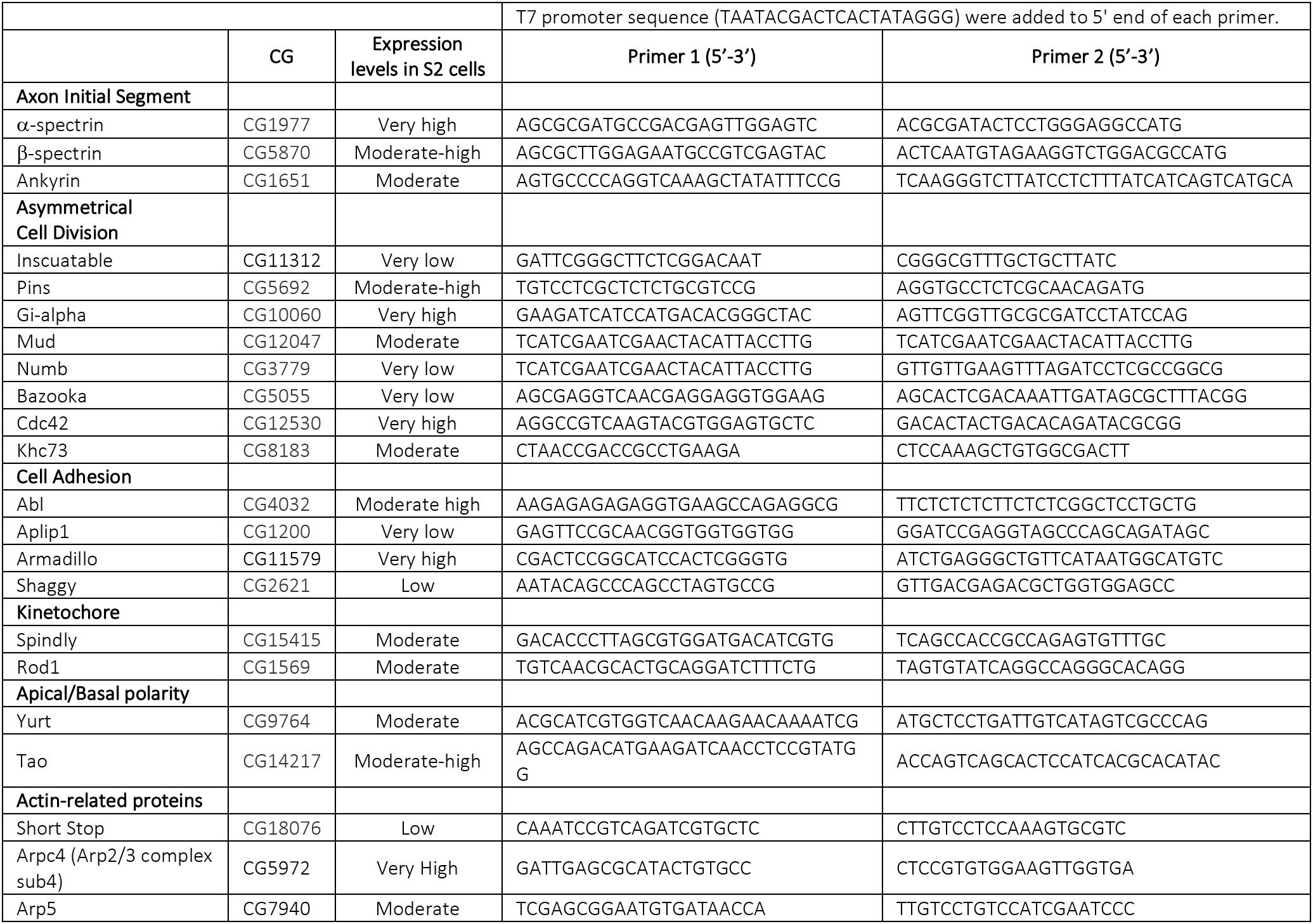
Genes tested in the candidate-based RNAi screen in S2 cells. Expression levels of these genes in S2 cells were obtained from (44).

